# Arc controls organ architecture through modulation of Crb and MyoII

**DOI:** 10.1101/2024.07.29.605562

**Authors:** Ji Hoon Kim, Rika Maruyama, Kwon Kim, Devin A. Vertrees, Parama Paul, Kyla Britson, Deborah J. Andrew

## Abstract

Precise orchestration of morphogenetic processes is required to generate organs that are optimally situated within the organism, and that are of the right size and shape to fit and maximize functionality. Here, we describe the role of Arc, a large apical membrane-associated PDZ domain-containing protein, that works through the apical determinant Crumb (Crb) to limit MyoII activity during tissue invagination in the forming Drosophila salivary gland (SG). We show that loss of Arc, attenuation of Crb function, as well as increased activation of non-muscle Myosin II (MyoII) leads to the simultaneous internalization of more precursor cells than normal. Consequently, mature SGs are significantly shorter with more cells surrounding the lumen at all positions along the tube. Correspondingly, overexpression of Arc or SG-specific knockdown of MyoII leads to the formation of longer SGs with fewer cells surrounding the lumen. We show that both Arc PDZ domains are required for Arc function and that they have distinct activities. Finally, we show that Arc facilitates Crb plasma membrane (PM) localization and suggests a model wherein PM-associated Crb stabilizes cellular junctions countering the destabilizing effects of apical medial and junctional pools of activated MyoII, thus limiting the number of primordial cells internalizing at any given time.

## Introduction

Epithelial tubular organs are essential for viability in all higher multicellular organisms. These organs, many of which are generated from polarized epithelial sheets, transport and exchange nutrients, wastes, and gases, and produce and secrete enzymes and hormones. Several life-threatening congenital conditions, such as esophageal atresia, pulmonary hypoplasia, and multiple forms of anorectal malformation, result from abnormal tube formation in the corresponding organs (Ioannides et al., 2010; Schittny, 2017; Wood & Levitt, 2018). Furthermore, around 90% of cancers are derived from epithelial tissues (Birchmeier et al., 1996; Hinck & Nathke, 2014). Therefore, investigating the molecular and cellular mechanisms underlying morphogenetic and homeostatic processes in epithelial tubular organs not only promotes our understanding of the biology of these essential organs but also carries crucial clinical implications.

Multiple model systems in various organisms have been used to study epithelial tubular organ development (Andrew & Ewald, 2010; Chung & Andrew, 2008; Lubarsky & Krasnow, 2003). Among them, salivary gland (SG) morphogenesis during Drosophila embryogenesis provides an excellent platform for uncovering the molecular mechanisms and cellular processes required for epithelial tube formation. SG cells are specified as a pair of two-dimensional sheets (SG placodes) located on the ventral side of the posterior embryonic head region flanking the ventral midline (**Fig. 1A,B**) (Chung et al., 2014; Girdler & Roper, 2014). Formation of tubes from these placodes is initiated by invagination of a small subsets of SG cells at dorsal posterior positions within the placodes (the invagination pit) (**Fig. 1I-N**) followed by the consecutive and orderly internalization of the remaining precursor cells to generate completely internalized, unbranched, elongated epithelial tubes (**Fig. 1C-H**) (Myat & Andrew, 2000b). During this process, SG cells undergo dynamic changes in position and shape such as cell intercalation and apical constriction, which facilitate movement toward and internalization through the invagination pit (Chung et al., 2017; Myat & Andrew, 2000b; Sanchez-Corrales et al., 2018).

**Figure 1.**
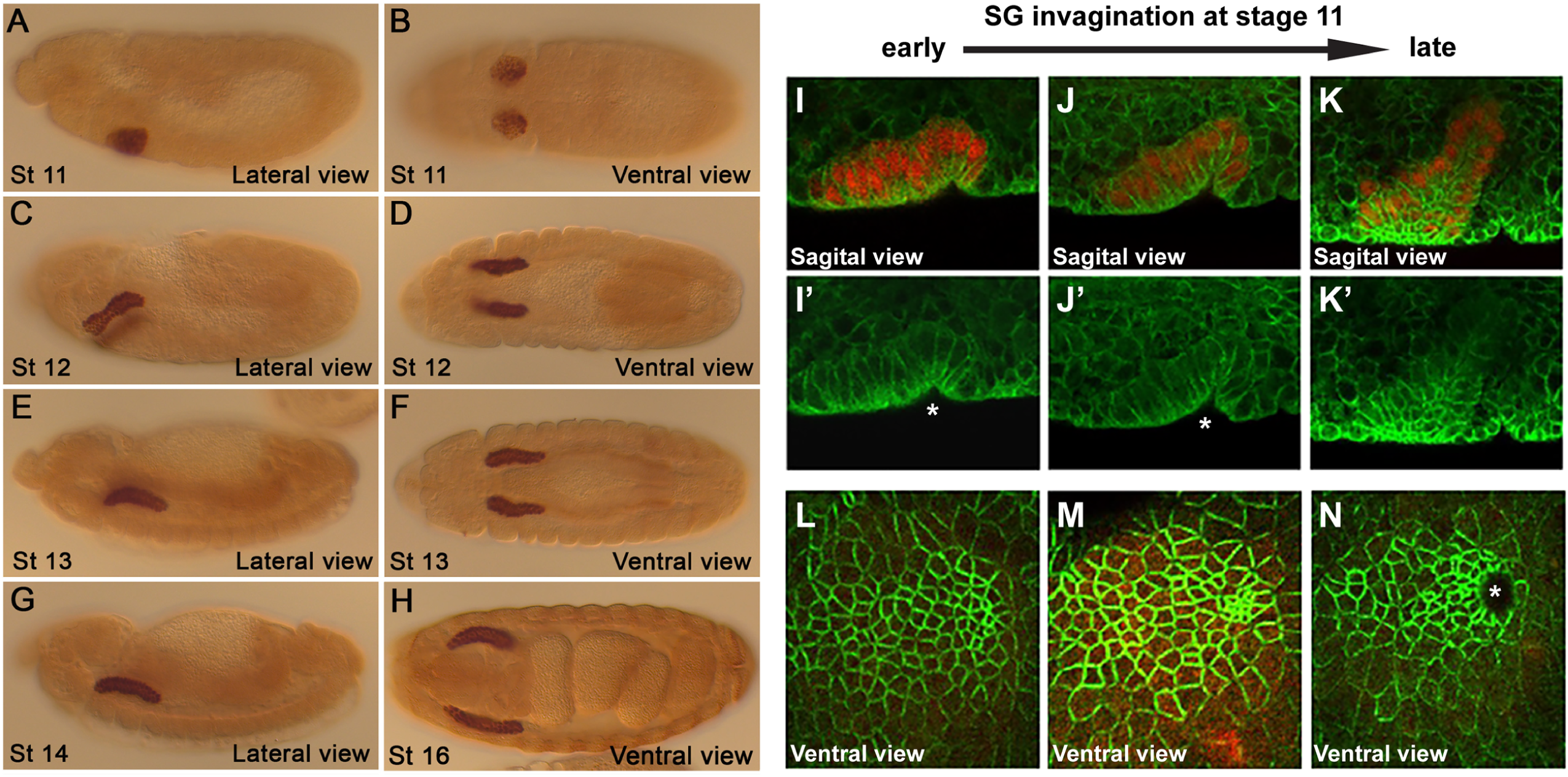
Salivary gland internalization begins in a dorsal apical domain with apical constriction. (A-H) detection of Sage nuclear protein in WT (Oregon R) embryos. Left panels show lateral views, right panels show ventral views. Salivary gland invagination begins in stage 11 (St. 11) and is largely complete by St. 13. (I-N) SG internalization begins with apical contriction of cells in a doral posterior position. WT embryos stained with nuclear Sage (red) and the adherens junction (AJ) protein E-Cadherin (ECad, green) (I, J, K, L-N) or just Ecad (I’, J’, K’). Asterisks mark the invagination pit.

A key molecular player driving SG morphogenesis is non-muscle myosin II (MyoII), a motor protein that exerts contractile force on actin filaments (actomyosin complex). As in other epithelial tissues forming tubular organs (Leptin, 1991; Sweeton et al., 1991), MyoII exhibits characteristic accumulation patterns on the apicomedial plane (apicomedial MyoII) as well as at cell-cell junctions (junctional MyoII) in SG cells (Röper, 2012). At the outer boundary of the SG placode, a contiguous myosin cable connected through multiple cells (supracellular MyoII cable) encompasses the entire placode (Röper, 2012). Several studies from our group and others have shed light on the roles of each MyoII structure during SG invagination (Booth et al., 2014; Chung et al., 2017; Sanchez-Corrales et al., 2018). For example, apicomedial MyoII drives apical constriction, which transforms the columnar SG cells into the wedge-shaped cells that facilitate internalization (Booth et al., 2014; Chung et al., 2017). Junctional MyoII promotes junctional shrinkage and extension during neighbor exchange between epithelial cells (Bertet et al., 2004; Curran et al., 2017; Levayer & Lecuit, 2013), including those of the SG (Sanchez-Corrales et al., 2018). A recent study proposes that the supracellular MyoII cable functions as a mechanical insulator protecting neighboring tissues from the morphogenetic changes in the SG placode (Ashour et al., 2023), although this structure could also contribute to the forces driving SG internalization or stabilizing the overall apical area occupied by the SG.

Despite of the central roles of MyoII in SG invagination, our current understanding about how MyoII activity and localization is regulated in SG cells is incomplete. Whereas G protein-coupled receptor (GPCR)-mediated activation of the Rho1 GTPase-Rho kinase (Rok) has been considered as a direct upstream signal to activate MyoII (Chung et al., 2017), how MyoII activity is spatially restricted and temporally coordinated for efficient cell invagination in the SG placode is largely unknown. A potential candidate that plays a role in tuning MyoII activity is the transmembrane protein Crumbs (Crb). Both *crb* mRNA and protein are elevated in the SG placode in comparison to surrounding epithelial cells implying its active role in SG invagination (Kerman et al., 2008; Röper, 2012; Chung et al., 2017). Crb is well known for its functions in establishing apicobasal polarity and mediating cell-cell adhesion through homophilic interactions of its extracellular domain. The short cytoplasmic domain of Crb contains functional motifs such as a FERM-binding motif and a PDZ-binding motif to recruit downstream effectors and transduce signals (Pocha & Knust, 2013; Thompson et al., 2013). Crb has been shown to either increase or suppress MyoII activity in different cellular contexts. In ectodermal cell development and neuroblast ingression, Crb recruits a Rho GTPase guanine exchange factor (GEF) Cyst to activate Rho1 and eventually MyoII (Silver et al., 2019; Simões et al., 2022). In aminoserosal cells, Crb represses MyoII activity to stabilize adherens junctions (Flores-Benitez & Knust, 2015). At the border of the SG placode, anisotropy of Crb between SG cells and surrounding epidermal cells induces the formation of the supracellular MyoII cable through accelerated membrane dissociation of Rho kinase by Crb (Sidor et al., 2020). Whereas these examples demonstrate the complicated relationship between Crb and MyoII, it is evident that Crb is a salient regulator of MyoII function in developing epithelial tissues. Therefore, how Crb affects MyoII activity during SG invagination and how Crb activity is regulated in SG cells are pivotal questions to understand fundamental principles underlying epithelial tubular morphogenesis.

In this study, we report that a novel PDZ domain-containing protein Arc affects overall tube dimensions by regulating Crb during SG invagination. Loss of *arc* results in shorter, stubbier SG tubes whereas excessive Arc dramatically elongates SG tubes. Arc colocalizes with Crb at the apical junctions in SG cells and maintains optimal Crb protein level in the SG placode by facilitating Crb delivery to the junctional membrane. Crb reduction in *arc* mutant SG cells abnormally increases MyoII activation, and overexpression of Arc or Crb disrupts MyoII accumulation, indicating that Arc and Crb play an inhibitory role in MyoII activation during SG invagination.

## Results

### *arc* functions downstream of Fkh to control tube geometry

Formation of the Drosophila salivary gland (SG) is governed by multiple early expressed transcription factors (Chung & Andrew, 2014). Among these factors, the winged-helix FoxA family protein Fork head (Fkh) plays a major role. Fkh expression begins at the earliest stages of SG tube formation (Weigel, Bellen, et al., 1989) and is sustained through tube migration and elongation (Weigel, Jurgens, et al., 1989). Indeed, *fkh* continues to be expressed throughout the lifespan of the larval gland (Cao et al., 2007) and is also expressed in the adult SG (FlyAtlas, Anatomical Expression Data). SG cells mutant for *fkh* fail to invaginate and remain on the embryo surface (Myat & Andrew, 2000a), suggesting that Fkh regulates expression of genes controlling the transformation of the two-dimensional SG placode into an internalized, elongated, three-dimensional epithelial tube.

Through whole mount in situ hybridization and microarray screens to discover SG Fkh targets (Maruyama et al., 2011), we identified *arc* (*CG6741*; abbreviated as ‘*a*’ in Flybase), a gene originally discovered based on the “arcing” phenotype of the adult wings in mutants (Bridges & Morgan, 1919). *arc* is expressed in multiple epithelia, including low level expression in the embryonic epidermis with elevated expression in the embryonic SG, foregut (FG), hindgut (HG), and Malpighian tubules (MT), and in both the eye and wing imaginal discs (Liu & Lengyel, 2000). *arc* mRNA is expressed in the SG from the earliest stages of SG invagination through later stages of tube elongation (**Fig. 2A, B**)*. arc* expression in the SG, HG, and MT is lost in *fkh* mutant embryos (Liu & Lengyel, 2000) (**Fig. 2C, D**), indicating that *arc* is a downstream transcriptional target of Fkh in multiple embryonic tubular epithelia.

**Figure 2.**
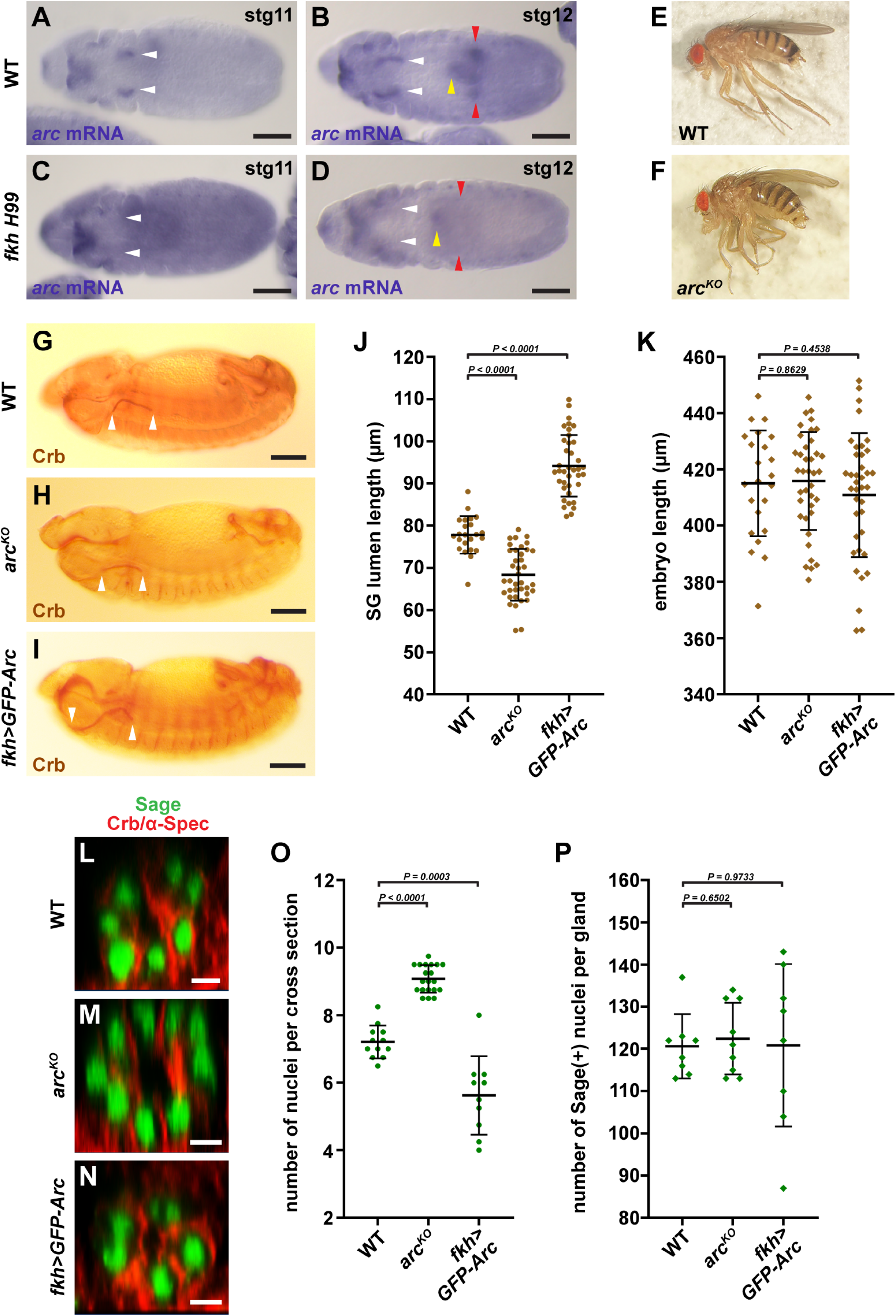
*arc* is regulated by Fkh and is required for proper SG morphogenesis. (A-D) detection of *arc* mRNA in WT (OregonR) (A, B) and *fkh H99* homozygous embryos (C, D) reveals Fkh-dependent expression of *arc* in the SG (white arrowheads), Malpighian tubules (red arrowheads), and hindgut (yellow arrowhead). *H99* is a genomic deficiency that deletes the pro-apoptotic genes and prevents the SG cell death associated with loss of *fkh*. Scale bars, 50 µm. (E, F) The wings of WT flies are planar whereas wings of *arc* nulls are downwardly curved in an ‘arc’ shape. (G-I) SG lumen length is altered with both loss of *arc* (H) and overexpression of *arc* using the *fkh-Gal4* SG driver (I). Arrowheads mark the proximal and distal ends of the SG secretory lumen. Scale bars, 50 µm. (J, K) Quantification of SG lumen length. Shorter SGs are observed with loss of *arc* and longer SGs are observed with excessive Arc (J); overall embryo length is unaffected by loss of *arc* or SG overexpression of *arc* (K). Each dot represents a SG. Error bars indicate standard deviation. (L-N) Cross-sectional views of SGs. Sage (green) labels SG nuclei. Crumbs (Crb) and α-spectrin (α-Spec) (red) mark cell boundaries. Scale bars, 5 µm. (O, P) Quantification of SG nuclei number. More nuclei are observed in cross section with loss of *arc* and fewer nuclei are observed in cross section with excessive Arc (O). Each dot indicates the mean value of nuclei counts from four optimally spaced cross-sections of a single SG. Total SG nuclei numbers are unaffected by *arc* loss or overexpression (P). Error bars indicate standard deviation.

To begin to study the role of *arc* in SG morphogenesis, we generated *arc* null alleles by replacing most of the coding region of *arc* with the *white+* eye color gene by homologous recombination (hereafter referred to as *arc^KO^*; **Fig. S1A, B**). *arc^KO^*flies are viable and demonstrate the same ‘arc’ wing defect as reported with the original *arc* mutant (Bridges & Morgan, 1919) and with a P-element insertion allele (*arc^k11011b^*) (**Fig. 2E, F**; (Liu & Lengyel, 2000)). *arc* mRNA was not detected in either of the two *arc^KO^* mutant lines we generated (**Fig. S1C**), confirming that both *arc^KO^* alleles are genetically null. Since the two KO alleles are molecularly identical, all subsequent analyses were carried out with *arc^KO788^*.

### arc affects SG tube dimensions

In *arc^KO^* mutant embryos (hereafter *arc^KO^*indicates both maternal and zygotic loss of *arc*), SGs fully invaginated, migrated and extended to form tubes (**Fig. 2G, H**); the length of SGs in *arc* null embryos (as measured by Crb-stained lumen length), however, was significantly shorter than in wild-type (WT) embryos (avg. SG lumen length WT 77.8 µm vs. *arc^KO^* 68.4 µm; **Fig. 2G, H, J**). Correspondingly, overexpression of Arc in the SG (*fkh-Gal4*-driven GFP-tagged Arc expression) resulted in longer SG tubes (avg. length WT 77.8 µm vs. *fkh>GFP-Arc* 94.19 µm; **Fig. 2G, I, J**). The changes in SG length with loss and overexpression of *arc* are not linked to changes in overall embryo length; WT, *arc^KO^*, and *fkh*>GFP-*arc* embryos were approximately the same length (avg. WT = 415.1 µm; *arc^KO^*= 415.9 µm; *fkh*>GFP-*arc =* 410.9 µm; **Fig. 2K**). The SG shortening with loss of *arc* and the lengthening observed with excessive Arc was consistently observed through all embryonic developmental stages (**Fig. S2A**). Furthermore, SG lumens were wider in *arc^KO^* than in WT during invagination (**Fig. S2C-E**). Taken together, these observations suggest that Arc affects SG tube dimensions from very early stages.

Once specified, WT SG cells cease proliferation and enter the process of polytenization (DNA replication without strand separation or nuclear or cell division) (Smith & Orr-Weaver, 1991). Likewise, there is no SG cell death during the entirety of embryogenesis and larval life in WT embryos (Cao et al., 2007; Myat & Andrew, 2000a). Thus, how the SG cells are arranged could impact overall tube dimensions. Shorter SG tubes may have more cells in circumference, whereas longer tubes may have fewer. To ask if this is the case in *arc* mutants, we counted the number of cells in cross-sections of late embryonic SG tubes. Indeed, the shorter SGs in *arc^KO^* embryos had more cells (avg. 9.08) in circumference than those in WT embryos (avg. 7.21) (**Fig. 2L, M, O**), whereas the longer SGs observed with overexpression of GFP-Arc had fewer cells in circumference than WT (avg. 5.63) (**Fig. 2N, O**). Since the overall cell numbers in the SGs of each genotype were approximately same (**Fig. 2P**), we conclude that the morphological changes observed with perturbations in *arc* expression result from defects in cell arrangement. Similar defects in tube dimensions were observed with *arc* loss in the hindgut, another epithelial tubular organ that expresses *arc*. As with the SG, hindgut length is significantly decreased with loss of *arc* (avg. hindgut lumen length WT 135.1 µm vs. *arc^KO^* 116.3 µm; **Fig. S3**). Taken together, these findings indicate that *arc* plays a key role in controlling the shape of epithelial organs during embryonic development by affecting cell arrangement.

### Arc colocalizes with Crb

To investigate how Arc affects cell arrangement during SG morphogenesis, we probed the localization of the endogenous Arc protein. Immunostaining with anti-Arc antibody combined with super-resolution confocal imaging revealed that Arc protein localizes to apical junctions and to punctate vesicular structures within the SG cells (**Fig. 3A, B**). Interestingly, junctional Arc often colocalized with junctional Crumbs (Crb), especially in regions of intense staining (**Fig. 3A-B”’**). The two proteins do not colocalize in the surrounding epidermal cells where Crb has a tight apical localization and Arc localizes along the lateral membrane (**Fig. 3B-B”’, arrow**). The short cytoplasmic domain of Crb contains an essential PDZ domain-binding motif (PBM) (Klose et al., 2013). The large Arc protein isoform expressed during embryogenesis (1329 amino acid residues) contains two PDZ domains in its C-terminal half allowing for potential direct interaction between Crb and Arc (**Fig. 3C**). To determine if one or the other PDZ domain of Arc is required for colocalization with Crb, we co-expressed Crb and different PDZ-deleted versions of Arc in cultured S2 cells (**Fig. 3D-G”**). Crb occasionally localized to sites of contact between S2 cells, which allowed us to examine localization of the co-expressed Arc proteins. We found that the first PDZ domain (PDZ1) is required for Arc colocalization with junctional Crb but the second PDZ domain (PDZ2) is not, as shown by lack of Arc accumulation to the Crb-enriched cell-cell contacts with Arc proteins that do not have PDZ1 (**Fig. 3D-G’’**). The PDZ1 domain is also required for apical confinement of junctional Arc protein in the SG. When overexpressed in the SG, PDZ1-deleted Arc proteins (Arc^ΔPDZ1^, Arc^ΔPDZ1/2^) were also observed in the lateral domain of SG cells where Crb is not found (**Fig. 3H-K”**). This suggests that the apical localization of Arc could be, at least in part, through direct physical interactions between Arc-PDZ1 and Crb-PBM. However, we failed to detect co-immunoprecipitation of Crb and Arc from extracts of cultured cells or embryos, suggesting that either the interaction between Crb and Arc is too transient or weak for this biochemical assay and/or that the colocalization of these two proteins requires additional factors. Intriguingly, when overexpressed ArcΔPDZ1 caused SG tube over-elongation similar to the full-length Arc, but ArcΔPDZ2 and ArcΔPDZ1/2 did not (**Fig. S4**). These results suggest that each PDZ domain has distinct molecular functions.

**Figure 3.**
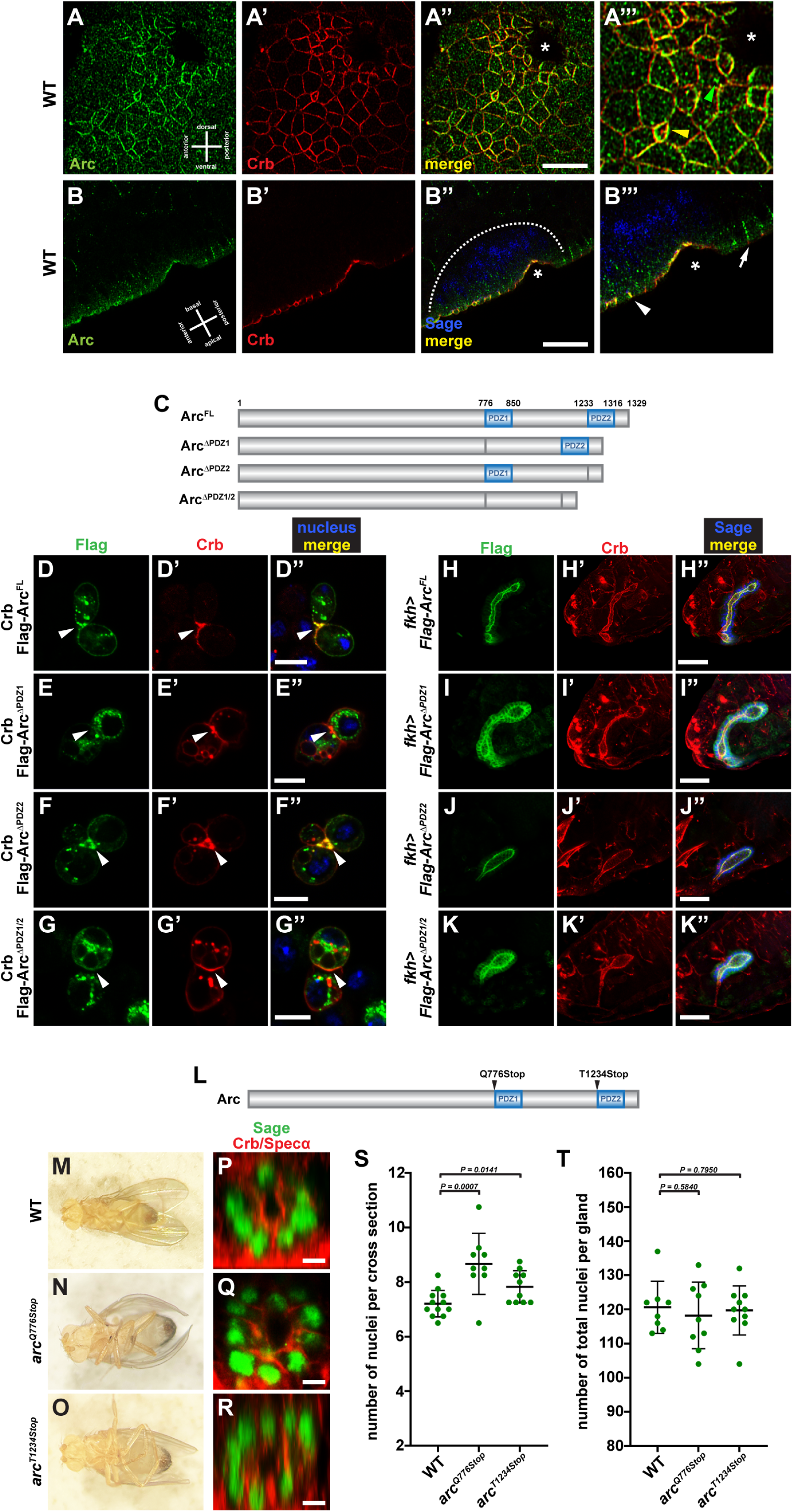
Arc protein is enriched at the cell-cell junctions in invaginating SG cells and co-localizes with Crb in a PDZ domain-dependent manner. (A-B’’) Immunofluorescence images of the SG placode in stage 11 embryos stained with α-Arc (green) and α-Crb (red). A-A’’’ show ventral view and B-B’’’ lateral view. Sage marks nuclei in SG cells (blue) encompassed with the dotted line (B’’). Note that Arc and Crb colocalize well in bicellular junctions when levels of both are high (yellow arrowhead in A’’’), but that they do not always colocalize, especially in the Arc punctate vesicular structures (green arrowhead in A’’’). Arc localization with Crb at apical junctions is specific to the SG (arrowhead in B’’’) and is not seen in the surrounding epidermis (arrow in B’’’). Asterisks mark the invagination pit. Scale bars, 10 µm. (C) Schematic drawing of Arc proteins with PDZ deletions. (D-G’’) Co-expression of Arc proteins with PDZ deletions with Crb in cultured S2 cells. Arrowheads indicate cell-cell contacts where Crb is accumulated. Full-length and PDZ2-deleted Flag-tagged Arc (green) (Arc^FL^ and Arc^ΔPDZ2^) co-localize with Crb (red) (D’’ and F’’) but PDZ1- and PDZ1/2-deleted Arc (green) (Arc^ΔPDZ1^ and Arc^ΔPDZ1/2^) do not (E’’ and G’’). Scale bars, 10 µm. (H-K’’) Localization of Flag-tagged Arc proteins (green) with PDZ deletions in SG tubes in stage 16 embryos. Deletion of the PDZ1 domain resulted in diffusion of Arc localization into the lateral domain (Arc^ΔPDZ1^ in I and Arc^ΔPDZ1/2^ in K), whereas localization of Arc^FL^ and Arc^ΔPDZ2^ was confined in apical domain (marked by Crb, red) (H and J). Scale bars, 50 µm. (L) The CRISPR/Cas9-mediated deletions in the *arc* gene. (M-O) The curved ‘arc’ wing phenotype in *arc* mutant flies. (P-R) Cross-sectional views of *arc* mutant SGs. Sage labels SG nuclei (green). Crb and α-spectrin (Specα) mark cell boundaries (red). Scale bars, 5 µm. (S, T) Quantification of SG nuclei number. More nuclei per cross section were seen in both *arc* mutant SGs compared to WT (S), whereas the total numbers of nuclei are similar (T). Each dot indicates a mean value of nuclei counts from four cross-sections in a SG. Error bars indicate standard deviation.

Based on the localization and overexpression results, we asked if the PDZ domains are necessary for Arc function in SG morphogenesis. Using CRISPR/Cas9 gene editing, we introduced nonsense mutations at the N-terminus of each PDZ domain in the *arc* gene locus to generate PDZ domain-deleted *arc* mutations (*arc^Q776Stop^* and *arc^T1234Stop^*) (**Fig. 3L**). Both *arc^Q776Stop^* and *arc^T1234Stop^* mutant flies have the arc wing defect demonstrating a general requirement for both PDZ domains in tissue morphogenesis (**Fig. 3M-O**). Removal of both PDZ domains (*arc^Q776Stop^*) resulted in approximately the same increases in cross-sectional nuclei number (avg. 8.67) as those observed in the *arc^KO^* mutant (avg. 9.08), indicating that the PDZ domains are essential for Arc function in SG morphogenesis. Deletion of the PDZ2 domain alone (*arc^T1234Stop^*) produced a milder defect (avg. 7.83), suggesting that both PDZ domains are required but may have different molecular functions in the SG (**Fig. 3P-T**). Taken together, Arc localizes with junctional Crb in SG cells in a PDZ1 domain-dependent manner and both PDZ domains are required for Arc function.

### Arc maintains Crb level in invaginating SG cells

Crb plays key roles in establishing and maintaining apicobasal polarity and transducing signals in many epithelial morphogenetic processes (Bulgakova & Knust, 2009; Tepass, 2012). During invagination, SG cells elevate Crb in a Fkh-dependent manner (Chung et al., 2017). Higher Crb level in SG cells than in surrounding epithelial cells is necessary for the formation of the supracellular MyoII cable around the SG placode (Ashour et al., 2023; Röper, 2012; Sidor et al., 2020). Furthermore, perturbations in Crb expression in the SG placode disrupt proper internalization of SG cells (Chung et al., 2017; Le & Chung, 2021), highlighting the importance of Crb regulation in tube morphogenesis. The colocalization of Arc with Crb prompted us to ask if Crb is affected in *arc* mutants. Strikingly, Crb levels in the SG placode were notably decreased with loss of *arc* (**Fig. 4A-B”**, **D**). Although excessive Arc did not significantly increase the overall amount of Crb, it did alter its distribution (**Fig. 4C-C”**, **D**, **6A-B’’’**).

**Figure 4.**
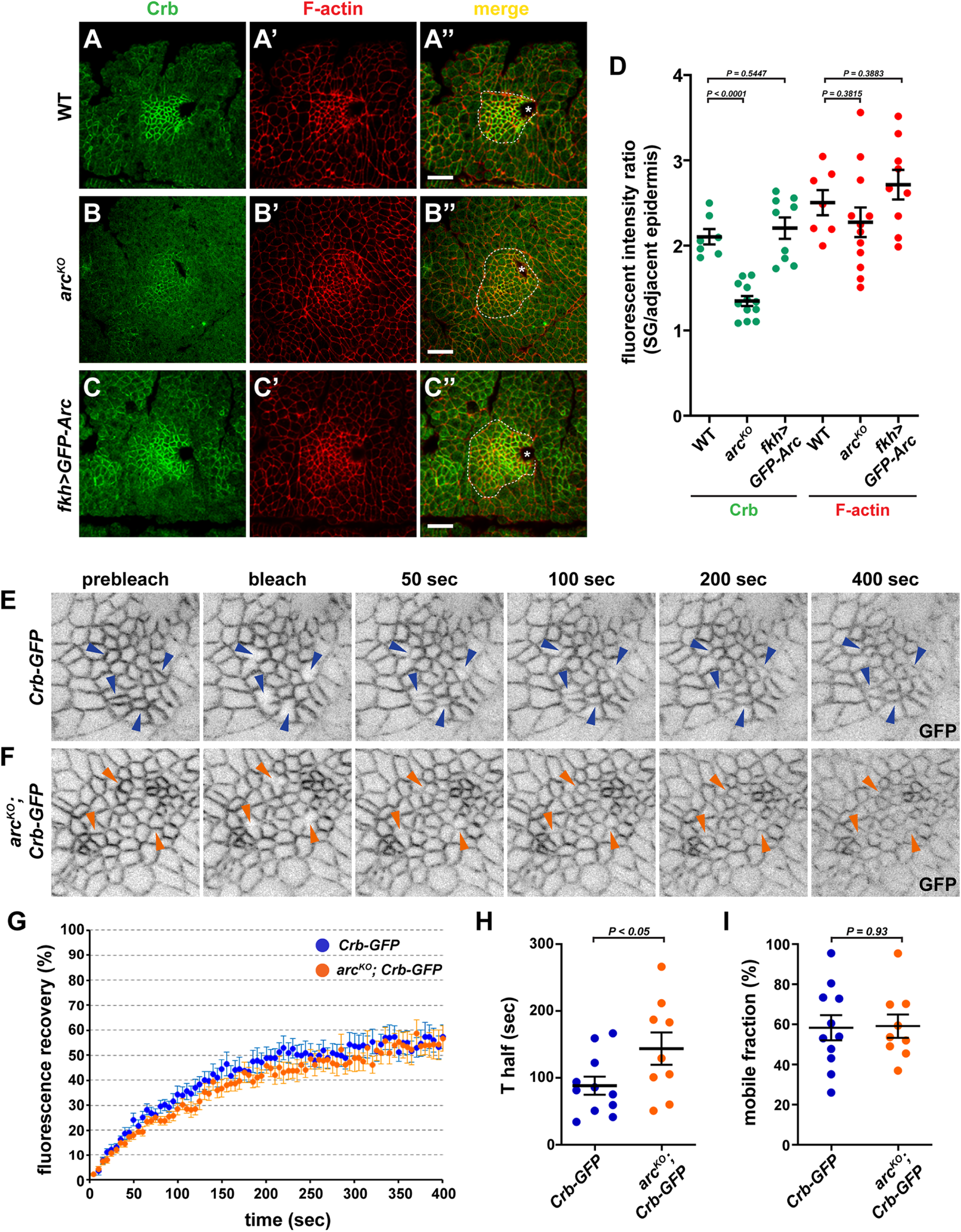
Arc maintains Crb levels in the SG placode. (A-C’’) Immunostaining of Crb protein (green) in invaginating SG placodes (encompassed with white dotted lines) in WT (A), *arc^KO^* (B), or expressing GFP-Arc (C). F-actin visualized by staining with phalloidin (red) (A’, B’, and C’). Asterisks mark the invagination pit. Scale bars, 10 µm. (D) The fluorescent intensities of Crb and F-actin in the SG placode normalized by those in adjacent epidermal area. Each dot represents a single SG placode. Error bars indicate standard error of mean. (E, F) Still images from FRAP time-lapse movies of GFP-tagged endogenous Crb in WT (E) and *arc^KO^* (F) embryos. Arrowheads indicate bleached cell junctions. (G-I) The recovery kinetics of the GFP fluorescence of Crb-GFP in WT (blue) and *arc^KO^* (orange) after photobleaching. Each dot represents a single junction. Error bars indicate standard error of mean.

To begin to understand how Arc affects Crb levels at SG cell junctions, we did fluorescent recovery after photobleaching (FRAP) to measure the exchange of Crb proteins on the cell membrane. In the wild-type placode, endogenous Crb tagged with GFP exhibited a dynamic behavior as shown by the significant recovery after photobleaching (avg. T_half_ = 88.23 sec, avg. mobile fraction = 58.33%). In the *arc^KO^* placode, Crb recovered to the same extent (avg. mobile fraction = 59.13%) as in wild type but at a significantly slower rate (avg. T_half_ 143.8 sec) (**Fig. 4E-I; Mov. S1**). This result suggests that Arc facilitates replenishment of Crb to cell-cell junctions in invaginating SG cells to maintain optimal Crb levels for proper SG morphogenesis.

### MyoII activity is altered in *arc* mutants

Non-muscle myosin II (MyoII) exerts contractile forces on the actin cytoskeleton to drive the changes in SG cell shape and arrangement during invagination (Booth et al., 2014; Chung et al., 2017; Sanchez-Corrales et al., 2018). We find that MyoII activity is critical for proper SG cell internalization as shown by the failure of SG cells to fully internalize with SG-specific nanobody-mediated depletion of the myosin light chain (Drosophila protein Spaghetti squash, Sqh) (**Fig. 5A-B’**). As in other epithelial cells undergoing morphogenetic changes (Martin et al., 2009; Simões et al., 2017), MyoII is concentrated in an apicomedial domain and in apical cell-cell junctions in invaginating SG cells (Booth et al., 2014; Chung et al., 2017; Röper, 2012). Interestingly, MyoII and Crb localize in a mutually exclusive pattern within the SG placode suggesting possible inhibitory interactions between the two proteins (**Fig. 5C**, **C’**). Indeed, Crb has been reported to regulate MyoII function either positively or negatively in different cellular contexts (Biehler et al., 2021; Flores-Benitez & Knust, 2015; Sidor et al., 2020; Silver et al., 2019; Simões et al., 2022), suggesting that Crb regulation by Arc in SG cells may impinge on MyoII activity during invagination.

**Figure 5.**
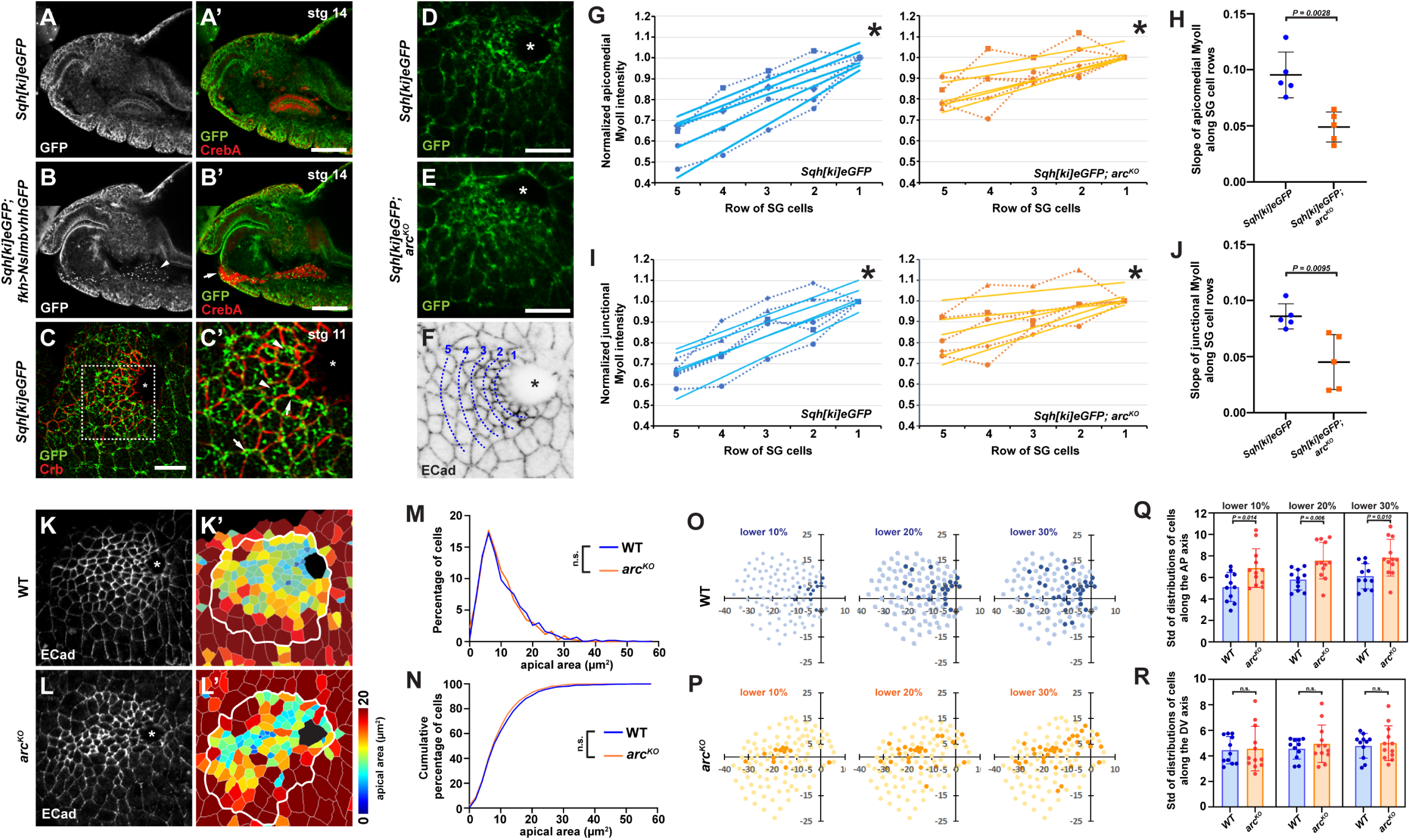
Arc affects MyoII activation and apical constriction in invaginating SG cells. (A-B’) The SG invagination defect caused by MyoII depletion. Expression of a GFP nanobody targeting the GFP-tagged endogenous Sqh resulted in a loss of MyoII (red) in its normal SG domains and localization to small aggregates (arrowhead in B) and frequent invagination failure (arrow in B’). Scale bars, 50 µm. (C-C’) MyoII localization visualized by anti-GFP immunostaining of the endogenous myosin light chain (Sqh) tagged with GFP (green). Crb marks cell-cell junctions (red). C’ is the enlarged view of the white box in C. Note that MyoII (green) is enriched at apicomedial domains (arrowheads) and cell-cell junctions (arrows) in the SG placode and that there is no overlap with junctional Crb (red). Scare bar, 10 µm. (D, E) Immunostaining of MyoII in WT (D) and *arc^KO^* SG placodes. Scale bars, 10 µm. (F) SG cell alignment in rows for MyoII quantification. Immunostaining of E-cadherin (ECad) marks cell boundaries. (G-J) The apicomedial (G) and junctional (I) MyoII intensity quantification in WT (blue) and *arc^KO^* (orange) SG cells. Five placodes were measured and plotted to show MyoII intensity per row. The slopes of MyoII gradient in each placode were shown in H and J. Note that MyoII levels decrease less in *arc^KO^* SG cells along with the distances from the invagination pit. (K-N) Apical area of SG cells in WT (K) and *arc^KO^*(L) placodes were determined using the Imaris Software. Asterisks in all pictures mark the invagination pit. (M) Percentage of cells with a given apical area and (N) cumulative percentage of cells of a given apical area in WT (blue) and *arc^KO^* (orange) reveals no differences. P values were calculated using the Mann-Whitney U test (percentage of cells) and using the Kolmogorov-Smirnov test (cumulative percentage of cells). (O-R) Plots of the distribution of cells with the smallest 10, 20, and 30% apical area in WT (O) and *arc^KO^* (P) SGs in a single example of each (K’, L’). The zero point on the X-Y axis is the center of the invagination pit. The standard deviations of distribution of cells of the smallest 10, 20, and 30 % smallest apical were plotted for WT and *arc^KO^* along the AP axis (Q) and the DV axis (R). Error bars are standard deviation. P values were calculated using the Student’s t-test.

To ask if Arc affects MyoII activity, we compared MyoII distribution throughout the SG placode in WT and *arc^KO^* embryos (**Fig. 5D-F**). In WT embryos, MyoII intensity is the strongest in both the apicomedial plane and in junctions in cells closest to the invagination pit; this intensity declines with increasing distance from the pit (**Fig. 5D, G, I**). Whereas *arc^KO^* SG cells present the same pattern in MyoII accumulation (highest at/near the invagination pit and lowest in the SG cells farthest from the pit), the steepness of the MyoII gradient throughout the placode is significantly reduced (**Fig. 5E, G-J**). Thus, without Arc, MyoII activity in the SG placode is abnormally elevated in cells several diameters away from the invagination pit. As expected from the abnormal MyoII activation, we found that apical constriction in *arc^KO^* SG cells is defective. Instead of a coordinated constriction pattern – smaller apical area in only SG cells close to the invagination pit (**Fig. 5K-K’**), *arc^KO^* SG cells quite distant from the invagination pit have abnormally small apical areas (**Fig. 5L-L’**). Surprisingly, the size distributions of apical areas of *arc^KO^* versus WT SG cells are no different (**Fig. 5M,N**); it is only the spatial distribution of cells with smaller apical areas that is changed with loss of *arc* (**Fig 5K’,L’**). Cells with the smallest apical area are distributed more anteriorly in *arc* mutants than in WT (**Fig. 5O-R**). These findings propound a molecular pathway wherein Arc promotes Crb localization. In turn, Crb attenuates MyoII activity to limit apical constriction to the subset of cells closest to the invagination pit, thus potentially limiting the number of SG cells that invaginate at any given time. Our findings also suggest that other factors affect the range distribution of apical area sizes in the invaginating SG.

### Arc overexpression phenocopies defects caused by excessive Crb

To test the model wherein Arc-dependent increases in Crb limit MyoII activity, we examined the localization of these proteins in SG placodes overexpressing Arc. Intriguingly, whereas Arc overexpression did not overtly increase the total amount of Crb in invaginating SG cells (**Fig. 3D**), it did alter Crb localization. In WT SGs, Crb localizes exclusively to apical junctions between cells, predominantly at bicellular junctions (**Fig. 6A-A’’’**). To our surprise, Crb also localized to diffuse punctate structures in/near the apical surface of Arc-overexpressing SG cells (**Fig. 6B-B’’’**). We detected similar ectopic Crb accumulation in diffuse punctate structures in or near the apical surface membrane in SG cells overexpressing Crb, although Crb levels are also increased at cellular junctions (**Fig. 6C-C’’’**). When overexpressed, Arc also accumulates to very high levels in apical-medial puncta in addition to its junctional localization, although Arc and Crb do not colocalize in these apical-medial puncta (**Fig. S5**). These observations indicate that excessive Arc (as well as excessive Crb) results in the delivery of Crb protein to cell membrane locations beyond its normal distributions.

**Figure 6.**
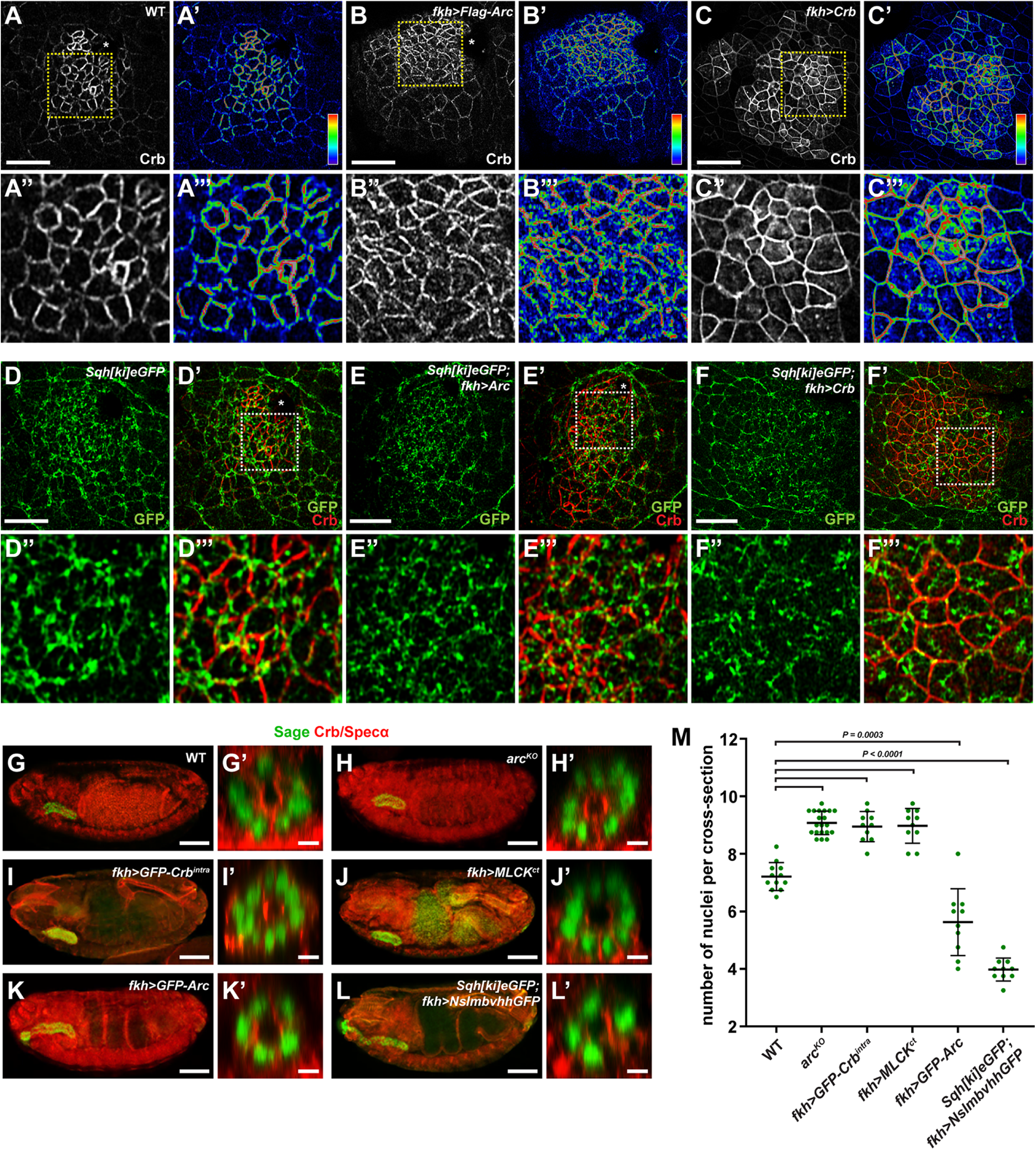
Arc overexpression alters Crb distribution, MyoII patterns and SG cell apical area. (A-C’’’) Immunostaining of Crb protein in WT (A), Arc-overexpressing (B), and Crb-overexpressing (C) SG placodes. Heatmaps represent the intensity of Crb staining (blue low, red high) (A’, B’, C’). The yellow boxes are magnified below. Note that the ectopic presence of Crb on the apicomedial plane in Arc- or Crb-overexpressing SG cells (B’’’, C’’’). (D-F’’’) Immunostaining of GFP-tagged endogenous Sqh in WT (D), Arc-overexpressing (E), and Crb-overexpressing (F) SG placodes (green). The white boxes are magnified below. Note the scattered MyoII distribution in small dots in the Arc- or Crb-overexpressing placode in contrast to the strongly aggregated pattern of MyoII in the WT placode. (G-J). Scale bars, 10 µm. Asterisks in all pictures mark the invagination pit. (G-L’) Immunostaining of SG nuclei with anti-Sage antibody (green). Crb and α-spectrin staining marks cell boundary (red). Representative cross-sectional images (G’-L’) are presented next to the whole embryo images (G-L). Note more nuclei with Arc loss (H’), Crb suppression (I’), and MyoII hyperactivation (J’) but fewer nuclei with Arc overexpression (K’) and MyoII depletion (L’). Scale bars, 50 µm (G-L) and 5 µm (G’-L’). (M) Quantification of SG nuclei number. Each dot indicates a mean value of nuclei numbers from four cross-sections in a SG. Error bars indicate standard deviation.

We next asked if MyoII is also affected by Arc or Crb overexpression since loss of *arc* resulted in decreases in Crb levels and increases in MyoII activation (**Fig. 4**, **5**). As described earlier, MyoII normally forms large trabeculate networks on the apicomedial plane and near apical junctions in SG placode cells close to the invagination pit (**Fig. 5C-C’**, **6D-D’’’**). In many SG placodes overexpressing Arc, MyoII formed smaller less contiguous aggregates (**Fig. 6E-E’’’**). A similar more exacerbated pattern of MyoII accumulation was observed in SG cells overexpressing Crb (**Fig. 6F-F’’’**). Although apicomedial MyoII was evident in SG cells overexpressing Crb, junctional MyoII was nearly gone. Thus, the ectopic accumulation of Crb, whether through overexpression of Arc or of Crb, disrupts MyoII activation, which is required for apical constriction. Indeed, we found that Arc- and Crb-overexpressing SG placode cells have similarly large apical areas and frequently failed to form an invagination pit at the right developmental stage (stage 11) (**Fig. 6B, C**). These findings support a model wherein Arc acts through Crb to attenuate MyoII activation.

### Functional relationship among Arc, Crb and MyoII

The changes in Crb and MyoII caused by Arc perturbation and the consequent defects in apical constriction prompted us to ask if altering Crb or MyoII activity affects SG tube morphology in a manner similar to loss or misexpression of Arc. We genetically manipulated the activities of Crb and MyoII specifically in the developing SG and probed SG tube dimensions by counting the nuclei number in cross-sections.

As reported by others (Tepass & Knust, 1990), we found that *crb* null mutant embryos exhibit severe developmental defects from very early stages, which makes examination of SG morphogenesis impossible. To circumvent this obstacle, we attempted to reduce Crb function specifically in SG cells by expressing a truncated Crb protein that contains only the transmembrane and intracellular cytoplasmic domains (Crb^intra^; (Klebes & Knust, 2000)). The Crb^intra^ protein localizes to the entire SG cell membrane (everywhere except where septate junctions form), unlike endogenous Crb, which is confined apically. Importantly, with Crb^intra^ expression, the endogenous Crb protein also localizes to the basolateral membrane thereby potentially reducing Crb levels and function in the apical domain (**Fig. S6**). Overexpression of Crb^intra^ resulted in shorter SGs with increased cross-sectional nuclei number (avg. 8.95), very similar to the SG phenotypes observed in *arc^KO^* embryos (avg. 9.08) (**Fig. 6H-I’, M**). Furthermore, hyperactivation of MyoII by overexpression of a myosin light chain kinase (MLCK; (Kim et al., 2002)) also generated phenotypes similar to loss of *arc*: shorter glands with more nuclei in cross-section (avg. 8.98) (**Fig. 6J, J’, M**). On the other hand, depletion of MyoII increased tube length and dramatically reduced the cross-sectional nuclei number (avg. 3.98), the same phenotypes as observed in embryos overexpressing Arc in the SG (avg. 5.63) (**Fig. 6K-M**). In summary, we have shown that loss of Arc, decreased Crb activity, as well as increased MyoII activity generates shorter SG tubes with more cells in circumference whereas gain of Arc and decreased MyoII activity results in abnormally long SGs with very few cells in circumference.

## Discussion

The SG of Drosophila is a great model for uncovering the molecules and mechanisms underpinning the morphogenetic processes of epithelial tube formation. Here we report on Arc, a large PDZ-domain containing cytosolic protein that functions downstream of Fkh to regulate the overall dimensions of at least two embryonic tubular organs. With loss of *arc*, we observe shorter SGs and shorter hindguts, with no corresponding change in overall embryo length. In the SG, where cell number is known to be invariant, we find that the length change in *arc* nulls reflects the arrangement of cells, with more SG cells in circumference and fewer SG cells along the length of the tube. Correspondingly, overexpression of Arc results in longer tubes with fewer cells in circumference. Our data support a model wherein Arc functions during early stages of SG internalization to promote Crb delivery to the apical plasma membrane. In turn, Crb modulates MyoII activity to control the number of cells that change shape and internalize at any one time, thus templating overall tube dimensions from the earliest stages of SG morphogenesis.

Pools of MyoII are critical for multiple aspects of morphogenesis. Apicomedial MyoII provides forces perpendicular to the apical junctions (AJ) to pull these junctions inward at points of contact (Martin et al., 2009). Junctional MyoII drives the corresponding shrinkage of apical junctions for apical constriction and for neighbor exchange through the formation of T1 junctions or multicellular rosettes (Bertet et al., 2004; Blankenship et al., 2006; Curran et al., 2017). Both apical constriction and neighbor exchange are critical to SG internalization, as these processes control which cells internalize in what order, as well as how many cells internalize at a given time, thus controlling overall tube geometry (Chung et al., 2017; Myat & Andrew, 2000a, 2000b, 2002; Sanchez-Corrales et al., 2018, 2021). Here, we show that Crb and MyoII exhibit complementary junctional localization within the SG placode prior to and during SG internalization: where Crb is high, MyoII is low, and vice versa (**Fig. 5C**). Related complementary patterns of Crb and MyoII accumulation are observed in SG cells at the boundary with the surrounding non-SG epithelium; junctional Crb is high where SG cells contact other SG cells but low where SG cells contact surrounding epithelial cells since, within the SG, Crb levels are much higher than in the surrounding epithelia (Chung et al., 2017; Kerman et al., 2008; Röper, 2012). A molecular pathway involving Crb recruitment of Par6, Cdc42 and Pak1 has been described for how Crb inhibits the membrane association of Rok (Sidor et al., 2020), a major activator of MyoII activity that acts through phosphorylation of the Myosin regulatory light chain (Yoneda et al., 2005). Thus, where Crb is high, junctional Rok and, consequently, junctional MyoII is low. Correspondingly, MyoII is high in the boundary junctions, leading to the formation of the supracellular MyoII cable surrounding the gland primordia (Röper, 2012). The mutual exclusivity between Crb and MyoII within and around the SG placode positions Crb as a key player in limiting MyoII activity.

Crb protein levels in epidermal cells are maintained by active endocytosis and recycling (Bajur et al., 2019). During apical constriction, actomyosin-driven contractions of the apical surface impact junctional Crb (and AJ associated proteins) by driving endocytosis and the disassembly of apical complexes (Bertet et al., 2004; Cavanaugh et al., 2020; Levayer & Lecuit, 2012). This turnover of apical complexes has been nicely demonstrated during neuroblast ingression, a process wherein individual isolated neuroblasts (NBs) undergo pulsatile apical contractions that serially decrease apical area through a ratcheting type process (Simões et al., 2022; Simões et al., 2017), much like occurs in mesodermal cells during gastrulation (Martin et al., 2009) and in SG cells during invagination (Booth et al., 2014; Chung et al., 2017). In NBs, apical area contraction and expansion during this ratcheting are accompanied by decreases and increases in junctional Crb levels, respectively. Importantly, the amplitude of apical constriction is dependent on the endocytosis of Crb and sorting of Crb for either degradation or recycling back to the junction. Loss of Retromer-mediated recycling function reduces both Crb-containing apical vesicles and Crb at apical junctions. Loss of Retromer-mediated recycling also accelerates apical shrinkage of NBs. This finding indicates that junctional Crb provides resistance to MyoII-mediated cell shape changes (Simões et al., 2022). Endocytosis of junctional Crb is normally limited by a PDZ-domain containing binding partner of Crb, Stardust (Sdt). With loss of *Sdt*, Crb is rapidly and completely endocytosed (Knust et al., 1993). Indeed, the completion of NB ingression – delamination of NB cells from the surrounding epithelia – requires the degradation of Sdt and the consequent depletion of junctional Crb (Simões et al., 2022). Taken together, these studies indicate that junctional Crb stabilizes apical junctions, functioning in a manner opposite to that of apicomedial and junctional MyoII, but in keeping with known functions of Crb in maintaining/expanding apical junctions in all primary epithelia (Tepass et al., 1990), including in the SG ((Chung & Andrew, 2014; Chung et al., 2017); this paper). This antagonistic relationship between Crb and MyoII in the SG and in the NB (Simões et al., 2022; Simões et al., 2017) highlights the importance of Crb regulation in epithelial morphogenesis.

Since Arc colocalizes with Crb at apical junctions and increases the rate of Crb recovery to these junctions based on our FRAP experiments, we propose that Arc facilitates the recycling of Crb to the apical surface to keep junctional levels of Crb appropriately high to counter MyoII activity. Arc-dependent Crb recycling could be mediated through direct interactions between PDZ1 of the Arc protein and the PDZ-domain binding motif (PBM) of Crb, which is located at the C-terminus of Crb’s small intracellular domain. Upon arrival at the apical surface, Arc may release Crb to allow for binding to Sdt, the PDZ-domain containing Crb-interacting protein that stabilizes junctional Crb (Tepass & Knust, 1993). This role for Arc would explain the relative depletion of junctional Crb in the absence of *arc* and the delivery of Crb to ectopic sites (apical medial membrane) when Arc is overexpressed. Our C-terminal deletions removing either one or both PDZ domains from the endogenous Arc protein indicates that both domains are required for WT Arc function. Whereas the PDZ1 domain is necessary to limit Arc to the apical domain of SG cells, the PDZ2 domain is required for elongating the SG tube when Arc is overexpressed, suggesting that this domain may function in the efficient delivery of Crb to its site of action.

Altogether, our data considered in light of published studies on Crb and MyoII in the SG as well as in other systems undergoing similar morphogenetic behaviors provide a working model for how Arc affects tube geometry (**Fig. 7**). SG internalization begins in a dorso-posterior position in the SG placode (Myat & Andrew, 2000a, 2000b; 2002), likely through the localized delivery of secreted Fog and/or the apical membrane localization of Fog’s corresponding GPCRs (Smog and TBD) (Le & Chung, 2021; Sanchez-Corrales et al., 2021; Vishwakarma et al., 2022). Upon receptor engagement, RhoGEF2, activated downstream of the receptor-associated G proteins, activates Rho1 GTPase and subsequently apicomedial Rok (Barrett et al., 1997; Levayer et al., 2011). In turn, localized Rok activates apicomedial Myosin, which drives apical constriction by pulling inward on the apical junctions (Booth et al., 2014; Chung et al., 2017). Based on the studies of NBs (Simões et al., 2017), we propose that apicomedial MyoII flows into the nearby apical junctions to shrink them through the endocytosis of Crb and AJ components (Bertet et al., 2004). The anisotropic flow of MyoII into the junctions results in the local endocytosis of Crb, creating domains of high MyoII, low Crb. This initial asymmetry is reinforced since Crb negatively regulates membrane associated Rok (Sidor et al., 2020), leading to the downregulation of activated MyoII in areas of higher Crb. Anisotropy in MyoII accumulation is important since SG cells must also rearrange with respect to one another so that the right number of SG cells internalize at any given time in the process (Sanchez-Corrales et al., 2021). Importantly, it is the variance in levels of apical junction associated MyoII that drives cell rearrangements (Curran et al., 2017). Thus, MyoII drives apical contractions and decreases in apical junction size. In concert with Crb, MyoII also drives directional cell rearrangement with Crb stabilizing contacts between neighboring cells and MyoII destabilizing these contacts.

**Figure 7.**
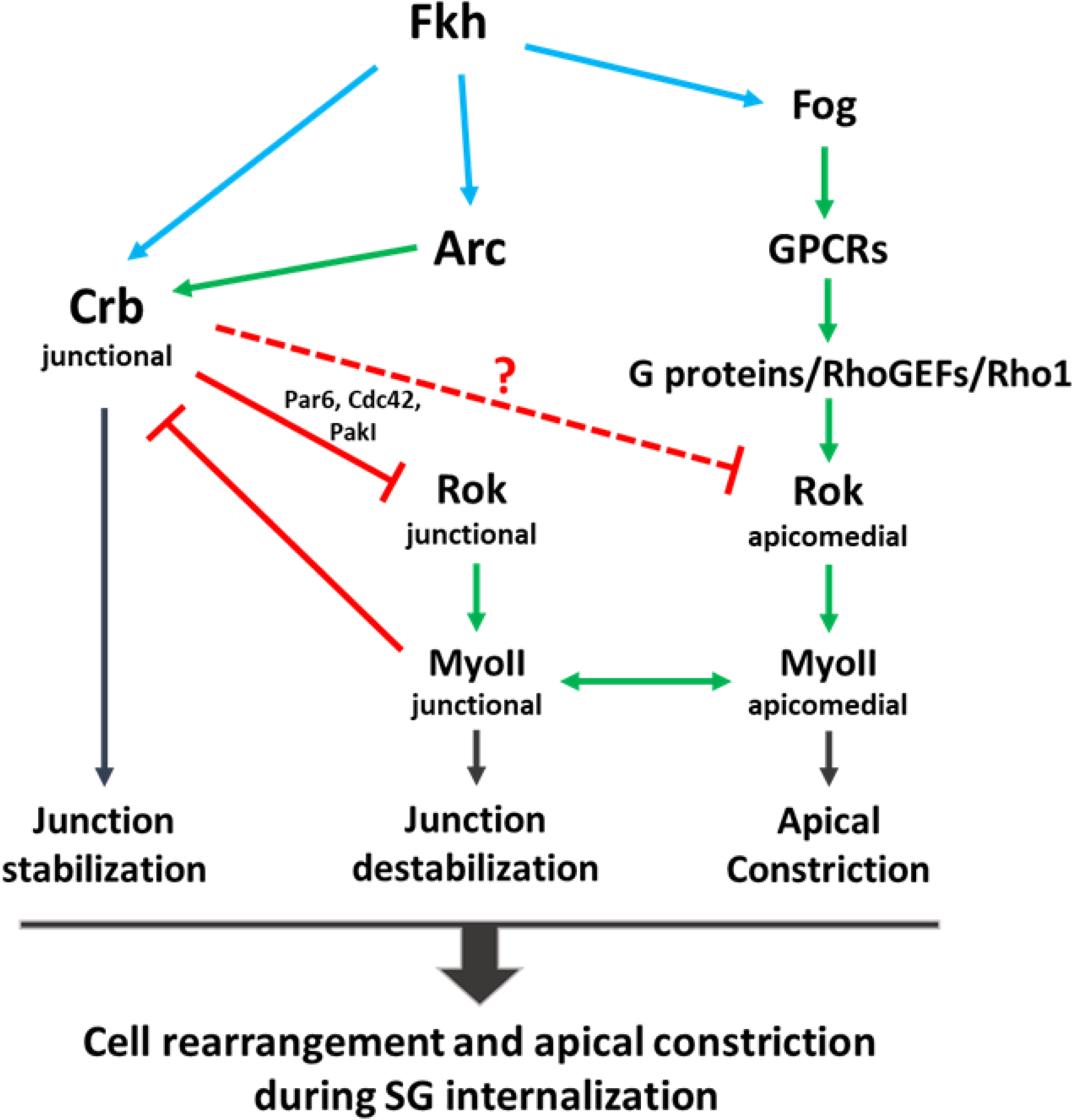
Model for Fkh regulation of SG internalization. Fkh activates expression of *arc*, *crb* and *fog* in the SG (this paper, Chung et al., 2017); each of these targets contributes to the dynamics of SG internalization ultimately determining final tube geometry. Fog functions as a ligand for the GPCR Smog and a likely to-be-determined additional SG-expressed GPCR (Vishwakarma et al., 2022). Fog binding to the SG GPCRs activates a signaling cascade that culminates in the activation of apical medial pools of Rok and MyoII, which pulls apical junctions inward (Chung et al., 2017). The apical-medial MyoII flows into junctions, creating anisotropic distributions of junctional MyoII, which locally counter the junction stabilizing effects of Crb, allowing for apical constriction and neighbor exchange. Acting in opposition of junctional MyoII accumulation, the complex that Crb forms with Par6, Cdc42 and Pak1 phosphorylates Rok, reducing its membrane association, thus preventing junctional MyoII accumulation in regions of high Crb. Arc functions to increase junctional plasma membrane (PM) pools of Crb and can also drive accumulation of apical medial Crb when overexpressed. Both the apical medial and junctional MyoII drive SG internalization, whereas plasma membrane associated Crb stabilizes junctions, counteracting the effects of Fog-dependent MyoII activation.

We propose that the increases in apical medial and junctional MyoII observed in cells several rows away from the invagination pit in *arc* mutants are a direct consequence of the reduced levels of Crb. We further propose that the consequent increases in MyoII support increases in both apical constriction and cell rearrangement, resulting in more SG cells internalizing during early stages of tube formation. We observe that overexpression of Arc results in increased levels of apicomedial Crb and with decreases in both apicomedial and junctional MyoII. In the apicomedial domain, Crb may interfere with Rok localization and/or activity. Reduced activation of MyoII in the apicomedial domain would consequently decrease junctional MyoII due to decreased flow. The consequence of these decreases in MyoII activation is that fewer cells undergo apical constriction and rearrangement, leading to the formation of tubes with fewer cells invaginating at any given time, with consequent longer tubes with a subset of cells that sometimes fail to internalize and remain on the embryo surface.

One interesting aspect of this study and related previous reports is that the transcription factor Fkh activates expression of two players in SG invagination that support mechanically opposing functions. Fkh activates *fog* expression in the SG (Chung et al., 2017). In turn, Fog activates MyoII through a GPCR signaling pathway that activates Rok in the apicomedial domain of invaginating SG cells (Chung et al., 2017). Rok-activated apicomedial MyoII drives apical constriction by pulling apical junctions inward. Independently, Fkh activates *arc* expression to ensure efficient delivery of Crb to apical junctions. At these junctions, Crb recruits the protein complex that phosphorylates Rok and decreases its membrane association, thus limiting Rok’s ability to activate junctional MyoII. These molecular pathways can explain the anisotropy in junctional MyoII and Crb distribution observed in SG cells. There, the two proteins have opposing activities, with MyoII driving junction disassembly and Crb stabilizing junctions. Thus, these Fkh downstream targets, through their opposing actions, fine tune the morphogenetic behaviors required to build epithelial tubes of proper dimensions (**Fig. 7**). We fully expect that the identification of additional Fkh target genes and investigation of their potential roles in modulating MyoII and Crb functions will lead to a deeper understanding of the molecular and cellular principles governing epithelial tubular architecture.

## Materials and Methods

### Fly genetics

The fly lines used in this study include: *Oregon R* (as wild type), *w^1118^* (as wild type), *fkh H99* (Myat & Andrew, 2000a), *fkh-Gal4* (Henderson & Andrew, 2000), *Crb::GFP-A* (Huang et al., 2009), *Sqh[ki]eGFP* (*Sqh-eGFP^29B^*; (Proag et al., 2019)), *UAS-GFP-Crb^intra^* (Pellikka & Tepass, 2017), *UAS-Crb*, *UAS-MLCK^ct^*, *UAS-Nslmb-vhhGFP* (Bloomington Drosophila Stock Center; https://bdsc.indiana.edu/). Two knock-out alleles of the *arc* gene (*arc^KO757^* and *arc^KO788^*) were generated by homologous recombination (Fig. S1; (Rong & Golic, 2001)). Both alleles are identical molecularly and phenotypically so are referred to as *arc^KO^* in the main text.

The PDZ domain deletion alleles, *arc^Q776Stop^* and *arc^T1234Stop^*, were generated by CRISPR/Cas9-mediated mutagenesis (Kanca et al., 2022). The individual alleles were identified by genomic DNA PCR amplification and sequencing. The UAS lines expressing full length or deleted versions of *arc* were generated by amplifying the *arc* ORF from cDNA GH21134 and cloning it into the pENTR-D/TOPO vector and then recombining the ORF into the Gateway vectors (http://flybase.org/reports/FBrf0179058) for expressing of either UAS-tagged (pTGW) or untagged (pTW) versions of the Arc protein. Flies carrying the UAS-constructs were then generated using P-element-mediated germline transformation (Rainbow Transgenic Flies Inc.).

### Molecular cloning of constructs for *arc* mutant and transgenic lines

To generate the knock-out construct for the *arc* gene, ∼4 kb of genomic DNA both upstream and downstream of the two largest coding exons of *arc* were amplified by PCR and cloned into the pW25 vector (Gong & Golic, 2004) on either side of the *white^+^* (*w^+^*) coding region (**Fig S1A**). Transgenic lines were generated and a line with an insertion of the construct on the 3^rd^ chromosome was subsequently crossed to a line expressing I-Sce1 under the control of a heat shock promoter. Progeny lines in which the *w^+^* marker had moved to the second chromosome (where *arc* localizes) were selected and analyzed by PCR for *w^+^* insertion into the endogenous *arc* gene (**Fig S1B**).

To generate deletions of the PDZ domains in the endogenous *arc* gene, we attempted the method of (Kanca et al., 2022) to cleanly delete each PDZ domain. This process includes synthesizing a DNA clone encoding two guide RNAs targeting the 5’ and 3’ regions of each PDZ domain and 200 bp immediately upstream and downstream of the corresponding PDZ domain, which should have yielded clean deletions of each PDZ domain. Although the guide RNAs worked well to cut the genomic DNA at the 5’ ends of each PDZ domain, homologous recombination did not occur with any of the lines that were generated. We did, however, succeed in generating lines by non-homologous end joining (NHEJ) that introduced premature stop codons deleting either one or both PDZ domains (*arc^Q776stop^*) or only PDZ-2 (*arc^T1234stop^*). To make the expression constructs of the *arc* gene (*UAS-Arc*, *UAS-GFP-Arc*, *UAS-Flag-Arc*, *UAS-Flag-Arc^ΔPDZ1^*, *UAS-Flag-Arc^ΔPDZ2^*, and *UAS-Flag-Arc^ΔPDZ1/2^*), the Gateway cloning system was used (http://flybase.org/reports/FBrf0179058). To specifically delete each PDZ domain from the *arc* ORF, the Takara In-Fusion cloning system was used following the manufacturer’s instruction (Takara).

### mRNA in situ hybridization and immunostaining in embryos

Whole-mount mRNA in situ hybridization was performed as described (Lehmann & Tautz, 1994) using an anti-sense digoxygenin-labeled *arc* RNA probe generated from cDNA GH21134. For immunostaining, embryos were fixed with 4% formaldehyde (Fisher Chemical) in PBS and devitellinized with methanol (Fisher Chemical). For α-Arc antibody staining, embryos were fixed by heat treatment as described (Liu & Lengyel, 2000; Peifer, 1993). For filamentous actin (F-actin) staining with phalloidin, the vitelline membrane was removed by hand after fixation with formaldehyde-saturated heptane (Sigma-Aldrich). The dilution ratios of primary antibodies follow: chicken α-GFP (1:500, Abcam), mouse α-Crumbs (1:10, DSHB), mouse α-alpha-spectrin (1:2, DSHB), mouse α-Arc (1;2000, a gift from J.A. Lengyel/V. Hartenstein), rabbit α-GFP (1:500, Molecular Probes), rabbit α-Flag (1:200, Proteintech), rabbit α-CrebA (1:100, (Fox et al., 2010)), rat α-E-Cadherin (1:20, DSHB), guinea pig α-Sage (1:200, (Fox et al., 2013)). Alexa Fluor-conjugated (Invitrogen) or biotin-conjugated (Jackson ImmunoResearch) secondary antibodies were used at 1:200. Alexa Fluor 488- or Alexa Fluor 568-conjugated phalloidin (ThermoFisher) were used at 1:200.

### Fluorescent recovery after photobleaching (FRAP)

For time lapse live imaging, embryos expressing GFP-tagged endogenous Crb (Crb::GFP-A; (Huang et al., 2009)) were collected and dechorionated in 50% bleach. After rinsing in water, embryos were placed onto double-sided tape (3M) on a microslide and covered in a layer of Halocarbon oil 700/27 (2:1; Sigma) with a coverslip. FRAP and time lapse image acquisition were carried out on an LSM 700 Meta confocal microscope (Zeiss). Regions of interest (ROIs) in a SG placode were manually selected and imaged in 5 frames to record the original fluorescent intensity (pre-bleach). Then the ROIs were quickly bleached to less than ∼20% of its original intensity (time point 0) and subsequently imaged with 30 second time intervals (post-bleach). The fluorescent intensities of the pre- and post-bleached ROI were measured using the image J program. The GraphPad Prism software (GraphPad Prism version 10.2.3 for Windows, GraphPad Software, www.graphpad.com) was used to determine the maximum recovery level (the percentage recovery to the pre-bleached level, mobile fraction) and the half-time of recovery (T_half_) in a kinetic curve fit with an exponential association equation.

### S2 cell culture and immunostaining

S2 cells were cultured in Schneider’s medium (Gibco) supplemented with 10% fetal bovine serum (FBS) (Gibco) and penicillin/streptomycin (Sigma). To express a protein of interest, the UAS vector encoding the protein and the ubiquitin-Gal4 construct were transfected with Effectene (Qiagen) according to the manufacturer’s protocol. For immunostaining, cells were fixed on a glass coverslip with 4% formaldehyde in PBS, washed in PBST (PBS with 0.1% Triton X-100) and PBSTB (PBST with 0.2% BSA) consecutively. The primary antibodies were used in PBSTB with the dilution ratios that follow: rabbit α-Flag (Proteintech, 1:1000) and mouse α-Crb (DSHB, 1:50). Secondary Alexa Fluor 488-, Alexa Fluor 568-, or Alexa Fluor 647-conjugated antibodies (Invitrogen) were used at 1:500. DAPI (ThermoFisher, 0.1 μg/ml) was used to stain nuclei.

### Image acquisition, analyses and quantification

DIC images of embryos were obtained on an Axiophot microscope (Zeiss) equipped with ProgRes CapturePro (Jenoptik). The whole adult fly images were taken by Iphone13 (Apple) on the Nikon SMZ800 microscope for magnification. Fluorescent images of embryos and S2 cells were obtained on an LSM 700 Meta confocal microscope or an LSM 880 Airyscan (Zeiss) equipped with Zen software (Zeiss). The obtained images were processed using Image J (http://imagej.nih.gov/ij/), Imaris (Bitplane) or Photoshop CS (Adobe).

To measure the lumen length of SG and hindgut, embryos were immunostained with α-Crb antibody. The lumen length was manually measured by tracking Crb signal in each tube using the Image J program. To count the number of SG nuclei in cross-sections, entire SGs immunostained with α-Sage, α-Crb, and α-αspectrin were imaged in multiple Z focal planes. The cross-sectional images were reconstructed from the Z focal images using the Image J program. The nuclei at four different cross-sections spaced optimally in one SG were counted and averaged to calculate the nuclei number for the SG.

To quantify Crb amount in SG placodes, embryos were immunostained with anti-Crb. Phalloidin was used to stain F-actin. Anti-Sage staining was used to define SG cells. The fluorescent intensity from an entire placode was measured and normalized by area To quantify MyoII intensity in apicomedial and junctional area respectively, embryos carrying the GFP-tagged endogenous *sqh* gene were immunostained with anti-GFP antibody. The fluorescent intensity from the GFP staining from apicomedial or junctional area in individual SG cells was manually measured using the Image J program.

The apical area of cells in a SG placode was determined by cell segmentation and calculation as described previously (Chung et al., 2017). Briefly, E-Cadherin images in SG placodes from one to three focal planes were integrated into one image by maximum intensity projection. The range of SG cells was determined by α-Sage staining. Cell segmentation based on the E-Cadherin signals, the subsequent apical area calculation and the colored map generation were performed by the Bitplane Imaris program version 10 (www.bitplane.com/imaris/imaris). All analyzed SG placodes were at the ‘deep invagination’ state in stage 11 in which SG cell under the invagination pit had internalized > 2 µm. Percentage distribution and cumulative percentage distribution of cells with a different apical area (bin width = 2) was performed using the GraphPad Prism software. For the spatial distribution analysis of SG cells, the x and y coordinates of each cell in a SG placode was determined using the Imaris software and used as the positions along the AP and DV axis from the center of the invagination pit at (0,0). Standard deviations for the positions along the AP or DV axis were calculated using the Microsoft Excel program and used to compare relative dispersion of SG cells.

## Supplemental Figure Legends

**Figure S1. Generation of *arc* null alleles by homologous recombination.** (A) *arc* knock-out alleles were created by replacing the two largest *arc* coding exons (3 and 4) with the *white^+^* (*w^+^*) eye color gene using homologous recombination. The PCR primers used to detect the replacement are marked with arrows in red font. LHA; left homology arm. RHA; right homology arm. (B) Insertion of the *w^+^* gene in two isolated alleles, *arc^KO757^* and *arc^KO788^* (both referred to as *arc^KO^* in the main text), was confirmed by PCR amplification. The combination of primers is in red font. The control is a PCR of the unrelated *Mvl* gene using Mvl-specific primers genomic template DNA from the *arc^KO757^* line that should yield a 2644 kb product. (C) In both *arc^KO757^* and *arc^KO788^* embryos, *arc* transcript is not detected by *arc* mRNA in situ hybridization. Scale bars, 50 µm.

**Figure S2. Arc affects SG tube dimension from early stages.** (A, B) Quantification of SG lumen length from stages 13-15. Loss of *arc* resulted in shorter tubes and Arc overexpression generated longer tubes at every stage. Each dot represents a SG. Error bars indicate standard deviation. (C-D’) Lumen width in the SG is wider in *arc* mutant embryos (D’) than in WT (C’). Arrowheads mark the width of SG lumen. Scale bars, 50 µm (C, D), 5 µm (C’, D’). (E) Quantitation of SG lumen width. Each dot indicates a mean value of three measurements at different locations along with a SG lumen. Error bars indicate standard deviation.

**Figure S3. The hindgut is shorter in *arc* null mutant embryos.** Immunostaining with anti-Crb antibody to visualize the hindgut in WT (A) and *arc^KO^* (B) stage 16 embryos. Scale bars, 50 µm. (C) Quantification of hindgut lumen length. Loss of *arc* resulted in shorter tubes. Each dot represents a hindgut. Error bars indicate standard deviation.

**Figure S4. PDZ2 is required for elongated SG tube phenotype by Arc overexpression.** (A-E) Immunostaining of SGs with anti-Crb (A; WT) or anti-Flag (B-E; overexpressing Flag-tagged Arc transgenes) antibodies in stage 16 embryos. Scale bars, 50 µm. (F) Quantification of SG lumen length. Only full length or PDZ1-deleted Arc extended SG lumen length when overexpressed. Each dot represents a SG. Error bars indicate standard deviation.

**Figure S5. Overexpressed Arc localizes apicomedially and junctionally in invaginating SG cells.** A series of focal planes from a SG placode immunostained for Flag-tagged Arc and Crb is shown. The depths of each plane are indicated from the most apical plane as 0 µm. When overexpressed, Arc is present on the apical membrane (arrowhead in A) as well as at the apical junctions (arrow in D). Scare bar, 10 µm. Asterisks mark the invagination pit.

**Figure S6. Overexpressed Crb^intra^ mislocalizes endogenous Crb proteins.** (A-A’’) anti-Crb immunostaining to visualize apical localization (red, arrowhead in A) of endogenous Crb in a WT stage 16 embryo. E-Cadherin (Ecad, green) marks apical junctions and Sage (blue) marks SG cell nuclei. (B-B’’) The GFP-tagged Crb^intra^ protein (that contains the transmembrane and cytoplasmic domains of Crb only) was expressed and visualized with anti-GFP immunostaining (green) in a stage 16 embryo (B’, B’’). Anti-Crb immunostaining (red) detects only endogenous Crb protein because the anti-Crb antibody recognizes the extracellular domain of Crb. Note that the endogenous Crb protein is present not only on the apical membrane (arrowhead) but also on the basolateral membrane (arrow) (B). Scare bars, 20 µm.

**Movie S1.** Fluorescent recovery after photobleaching (FRAP) time-lapse movies of GFP-tagged endogenous Crb in WT and *arc^KO^* embryos. Time 00:00 indicates the point of photobleaching. The bleached junctions are marked with arrowheads.

**Table S1.** PCR primer sequences used in this study.

## Author conflict statement

The authors declare that they have no conflict of interest.

